# Site-specific effects of online rTMS during a working memory task in healthy older adults

**DOI:** 10.1101/642983

**Authors:** Lysianne Beynel, Simon W. Davis, Courtney A. Crowell, Moritz Dannhauer, Wesley Lim, Hannah Palmer, Susan A. Hilbig, Alexandra Brito, Connor Hile, Bruce Luber, Sarah H. Lisanby, Angel V. Peterchev, Roberto Cabeza, Lawrence G. Appelbaum

## Abstract

The process of manipulating information within working memory (WM) is central to many cognitive functions, but also declines rapidly in old age. Given the importance of WM manipulation for maintaining healthy cognition, improving this process could markedly enhance health-span in older adults. The current pre-registered study tested the potential of online repetitive transcranial magnetic stimulation (rTMS) to enhance WM manipulation in healthy elderly adults. Online 5Hz rTMS was applied over the left lateral parietal cortex of 15 subjects to test the hypothesis that active rTMS would significantly improve performance compared to sham stimulation, and that these effects would be most pronounced in conditions with the highest cognitive demand. rTMS was applied while participants performed a delayed-response alphabetization task with two individually-titrated levels of difficulty. Sham stimulation was applied using an electrical sham coil that produced similar clicking sounds and somatosensory sensation as active stimulation but induced negligible effects on the brain. A stimulation site in left lateral parietal cortex was identified from fMRI activation maps and was targeted using individualized electric field modeling, stereotactic neuronavigation, and real-time robotic positioning, allowing optimal coil placement during the stimulation. Contrary to the *a priori* hypothesis, active rTMS significantly decreased accuracy relative to sham, and only in the hardest difficulty level. These results, therefore, demonstrate engagement of cortical WM processing, but not the anticipated facilitation, and provide a prescription for future studies that may attempt to enhance memory through application of different stimulation parameters.

**Highlights:**

- This study is one of the first attempts to enhance WM manipulation with online rTMS
- Online 5Hz rTMS and sham were applied over the left parietal cortex of older adults
- Individualized fMRI and electric field modeling were used to optimize targeting
- Contrary to expectations, rTMS disrupted working memory manipulation abilities
- This demonstrates that parietal cortex is involved in WM and modifiable with rTMS

## 1. Introduction

Working memory (WM), the capacity to maintain and manipulate information in a temporary mental buffer, is central to many aspects of human cognition. Indeed, through the interface between long-term memories and moment-to-moment information available in the environment, WM allows humans to organize relevant information in order to carry out successful goal-directed behaviors (Baddeley, 1998). As such, WM capacity is intrinsic to many daily activities such as reading, performing arithmetic, and keeping track of ideas during a conversation (Pasula et al., 2018; Kane et al., 2007). Both WM capacity and, more particularly, the ability to manipulate content that is held in WM declines with age (Cappell, Gmeindl, & Reuter-Lorenz, 2010; Kirova et al., 2015). Therefore, different approaches have been proposed to prevent this decline.

Brain stimulation techniques, such as repetitive transcranial magnetic stimulation (rTMS) in particular, have gained increased attention as a means to enhance WM and slow age-related impairments. rTMS uses brief, high intensity magnetic fields to depolarize neurons underneath a magnetic coil. When applied over a brain region that helps support a specific cognitive function, rTMS has the potential to modulate related behavior. In many such studies, rTMS applied *online*, i.e., during the performance of a task, has been shown to interfere with ongoing cognitive processes, thus impairing behavioral performance (see Walsh, & Pascual-Leone 2003 for a review). In other cases, however, studies have reported performance enhancement when applying online rTMS (see Luber & Lisanby, 2014 for a review). For example, stimulation of the parietal cortex during WM maintenance tasks has resulted in significant decreases in reaction times or accuracy improvement (Luber et al., 2007; Hamidi et al., 2008). These contrasting results suggest that online rTMS may affect performance in a manner that is specific to the ongoing process and the spatio-temporal parameters of stimulation, for example by modulating endogenous task-related oscillatory dynamics (Johnson, Hamidi, & Postle, 2010).

The application of neuromodulation in older adults presents unique challenges to cognitive neuroscience studies seeking to modulate cognitive performance. Aging has been associated with a relative decrease in the excitability of intracortical inhibitory circuits (Hortobagyi et al., 2006), and is also associated with a steady linear decline in cortical thickness (Salat et al., 2004) and cortical volume (Hutton et al., 2009), suggesting an important correction for the standards of a field largely based on younger adults. While linear adjustments of stimulation amplitude according to the distance from scalp to cortex (e.g., Stokes et al., 2007) may provide some correction for the systemic differences between older and younger adult brains, such linear adjustments ignore the multilinear changes in neuroanatomy associated with aging, and are fundamentally indirect estimates of how the TMS-induced electric field spreads across the cortical surface. There is reliable evidence that once these factors are controlled, the motor-evoked response to TMS does not differ significantly across the lifespan (Oliviero et al., 2006), suggesting that E-field modeling is a necessary component to any TMS study seeking to make normative inferences on the relationship between TMS stimulation, the selection of a specific stimulation site, and the engagement with underlying cortical oscillations associated with a particular cognitive function at that site.

Given the important role that theta oscillations play in memory processes (Roux, & Ulhas, 2014), our group has recently undertaken a series of studies aimed at testing the effects of 5Hz rTMS to facilitate the manipulation of information in WM. In one such study (Beynel et al., 2019), 5Hz rTMS was applied over the left dorso-lateral prefrontal cortex (DLPFC) while subjects performed a delayed-response alphabetization task (DRAT). During the DRAT, participants were briefly presented with an array of letters and asked to mentally arrange these letters into alphabetical order during a delay period, after they had been removed from view. Results revealed that both younger and older adults showed enhanced accuracy on the DRAT with active rTMS, compared to electrical sham stimulation, with these enhancements observed only in the most difficult conditions of the task. While this result was promising, the effect size remained modest and produced only a 4% improvement in memory recall. Extant theories of WM function suggest that while the DLPFC may play a role in mediating the online attentional processes associated with successful WM function, WM information is more centrally processed and stored in the lateral parietal cortex (Postle, 2006; Koenigs et al., 2009); thus, one potential way for optimizing rTMS neuro-enhancement protocol would be to target this more central location in the fronto-parietal WM network.

The goal of the present study was therefore to attempt to increase the potency of online rTMS for enhancing WM manipulation by stimulating another cortical area involved in this task. Because results from our previous fMRI study indicated that manipulation of information in the DRAT produced the greatest activation in the superior parietal lobule (Davis et al., 2018), fMRI-BOLD activity in the left lateral parietal cortex was used to derive the target for rTMS in the current study. This target was defined on an individual basis to determine TMS coil location (projected to scalp surface) and orientation (related to nearest sulci wall) by information derived from fMRI activation and brain anatomy (MRI). Electric field (E-field) modeling was then used to select the individual TMS amplitude to induce approximately the same E-field strength in the target region across subjects. This computational dosing method accounts for the individual head anatomy and was deployed in an effort to minimize individual variability of the response. Based on this experimental design, it was hypothesized that active rTMS would significantly enhance WM manipulation performance, and that this effect would be most pronounced in the most difficult task conditions, consistent with the view that cognitive performance is most vulnerable to neuromodulation under the most demanding conditions (Muggleton et al., 2003; Luber et al., 2007; Beck and Hallett, 2010)).

## 2. Material and Methods

### 2.1. Participants

Thirty-nine healthy subjects (60–80 years old) were recruited into this single-blind randomized within-subject controlled trial, which was pre-registered on ClinicalTrials.gov (NCT02767323). All participants provided written informed consent, which was approved by the Duke University Institutional Review Board (#Pro00065334). Participants were excluded if they had any contraindication to TMS or MRI, current or past psychiatric disorders or neurological disease (n=1), or a total scaled score lower than eight on the Dementia Rating Scale-2 (Jurica, Leitten, & Mattis, 2001) (n=1). Participants were also excluded if they tested positive on a urine drug screen (n=2) performed poorly on the WM task during the initial visit (n=11), or experienced non-compliance, for example by responding in the task with random keys presses (n=3). According to these criteria, 18 participants were excluded, along with another 6 participants who withdrew participation for unspecified reasons. Fifteen subjects completed the full protocol (4 females, mean age: 66.13 ± 5.5 years old, mean years of education: 17.33 ± 1.79 years). Participants had normal, or corrected-to-normal, vision and were native English speakers. Participants were compensated $20/hour for their efforts with a $100 bonus if they completed all study activities.

### 2.2. Experimental Protocol

Participants were scheduled for 6 sessions: a consenting visit including screening, resting motor threshold assessment and practice at the delayed-response alphabetization task, followed by an MRI visit and four rTMS visits (Figure 1). The following sections give a brief overview of the methods, and additional information can be found in Beynel et al. (2019).

**Figure 1:**
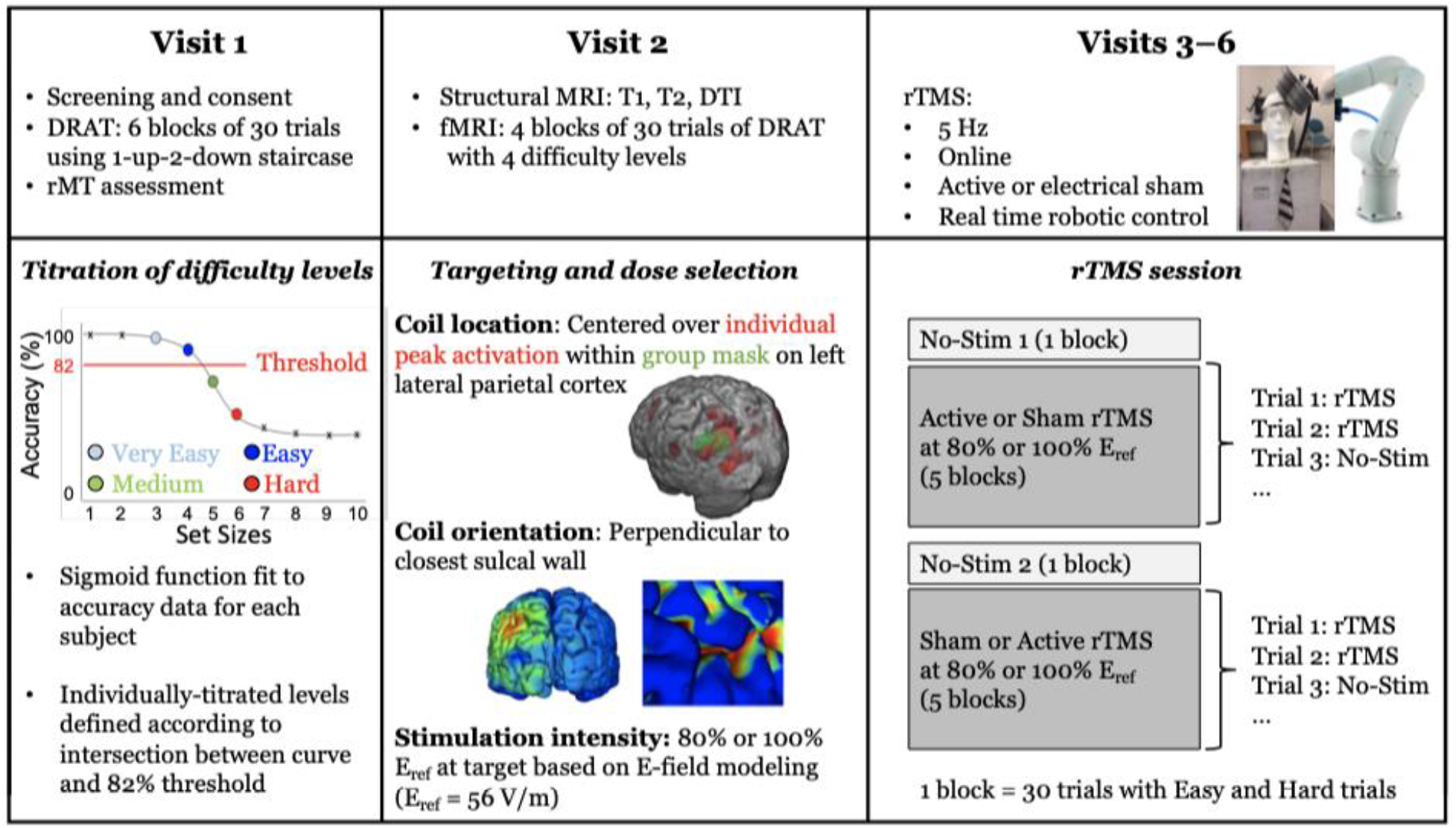
Illustration of the full experimental protocol across 6 visits. For visit 1 and 2, data processing procedures are illustrated in the bottom panel. For visits 3–6 the bottom panel illustrates the experimental procedure in greater detail.

### 2.3. Delayed-Response Alphabetization Task (DRAT)

On each trial of the DRAT an array of letters was presented on the screen for 3 seconds, followed by a 5-seconds delay period during which the participants were asked to mentally reorganize the letters into alphabetical order (Figure 2). After the delay period a letter with a number above it appeared on the screen for 4 seconds and participants were asked to report via key press whether the letter was (1) not in the original set, (2) in the original set and the number matched the serial position of the letter once the sequence was alphabetized, or if (3) in the original set but the number did not match the serial position of the letter once alphabetized. These conditions are referred to as New, Valid, and Invalid, respectively.

**Figure 2:**
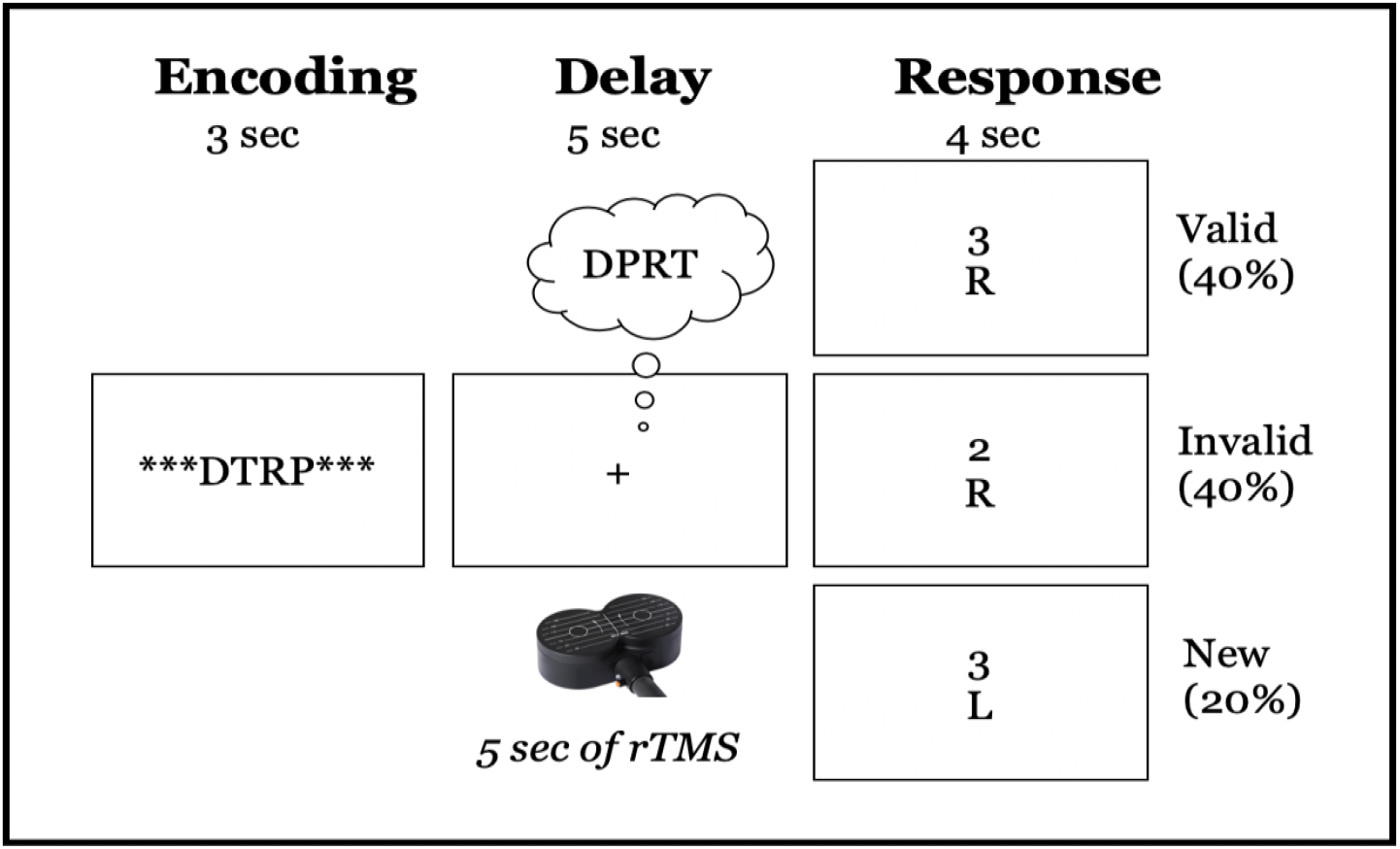
Schematic illustration of DRAT task. Top row shows an array of 4 letters participants are to encode, that is presented for 3-seconds. Row two shows the 5-second delay period, during which participants had to maintain and reorganize the letters into alphabetical order. Row three shows examples of the three possible responses: “New”: the letter was not in the original array; “Valid”: the letter was in the array and the number represented the correct position in the alphabetical order; “Invalid”: the letter was in the array but the number did not match the correct serial position when alphabetized.

During the first visit, participants performed the DRAT using a staircase procedure to establish individualized difficulty levels. Four individualized difficulty levels were defined according to the intersection between a sigmoid curve, fit to the data, and an 82% accuracy threshold. The two set sizes below this intersection were defined as Very Easy and Easy, and the two levels above it were defined as Medium and Hard. If the intersection between the curve and the threshold was lower than four, participants were considered poor performers and excluded from the study (n=11). While all four difficulty levels were used for the subsequent imaging visit, only the Easy and Hard levels were used during the TMS visits.

### 2.4. Targeting Approach

During the second visit, subjects participated in an MRI scanning (General Electric MRI Scanner, B_0_ field strength = 3 Tesla) during which structural-T1-weighted (echo-planar sequence: Voxel Size= 1 mm^3^, TR= 7.148 ms, TE = 2.704 ms, Flip Angle = 12°, FOV = 256 mm^2^, Bandwidth = 127.8 Hz/Pixel), T2-weighted (echo-planar sequence with fat saturation: Voxel Size = 0.9375 × 0.9375 × 2.0 mm^3^, TR = 4 s, TE = 77.23 ms, Flip Angle = 111°, FOV = 240 mm^2^, Bandwidth = 129.1 Hz/Pixel), and diffusion weighted scans (Single-shot echo-planar: Voxel Size = 2 mm^3^, TR = 17 s, TE = 91.4 ms, Flip Angle = 90°, FOV = 256 mm^2^, Bandwidth = 127.8 Hz/Pixel, Matrix size = 128^2^, B-value = 2000 s/mm^2^, Diffusion directions = 26) were obtained. Functional acquisitions (EPI-sequence: Voxel Size = 3.4375 × 3.4375 × 3.99 mm^3^, TR = 2 s, TE = 25 ms, Flip Angle= 90°, FOV = 220 mm^2^, Bandwidth = 127.7 Hz/Pixel) were also acquired as participants performed the DRAT in the scanner. After preprocessing the images, functional data were analyzed using a general linear model (GLM) in which trial events were convolved with a double-gamma hemodynamic response function. The GLM examined BOLD responses during trials where the correct response was chosen in the behavioral task. Separate events were modeled for the array presentation (3-second duration), the delay period (5-second duration), and the response period (trial-specific RT duration). All incorrect and non-response trials were modeled identically, but separately, and were not considered in the results.

The stimulation target was individually defined as the peak activation within the left lateral parietal cortex associated with a parametric increase in task difficulty during the delay period of the DRAT. According to the results obtained in our previous study (Davis et al., 2018), both Set Size (the number of letters in an array) and Sorting Steps (the minimum number of operations required to alphabetize the array) contribute to the difficulty of an individual trial. Therefore, to obtain a more accurate representation of increases in DRAT difficulty a parametric delay-period regressor, defined by the interaction between Set Size and Sorting Steps was used estimate task difficulty. This parametric regressor was orthogonalized to the non-parametric delay-period regressor. At the first level, functional data were analyzed as individual runs. Second-level analyses combined data across runs for each subject using a fixed-effects model. This processing allowed for the definition of the stimulation target on individualized statistical maps that predicted the parametric increase in BOLD activity associated with increasing task difficulty, if the peak activation reached a z-statistic value > 2; or alternatively on the nonparametric delay-period map if the peak did not reach this significance threshold.

To constrain the stimulation target within the left lateral parietal cortex, a mask obtained from the group activation of 22 older adults who participated in our previous study was used (Beynel et al., 2019). The mask was defined as the overlap between the parametric interaction between Set Size and Sorting Steps (at z > 1) and the non-parametric delay period activity (at z>1), therefore reflecting cortical regions that were generally activated by the task, but also were specific to difficulty increase. The individual activation was then transformed back into subject space and the peak activation within this mask was selected as the *TMS target* in the neuronavigation system (BrainSight, Rogue Research, Canada).

To define the coil orientation, the coordinates from the *TMS target* were projected onto the scalp surface using a nearest neighbor approach, and then projected slightly outwards to account for the subject’s hair thickness (**Supplementary, Figure S1**). Hair thickness was measured on each participant during the screening visit, using a depth gauge (Digital Tread Depth Gauge, Audew, Hong Kong, resolution: 0.01 mm) installed on a custom-made plastic base placed over the center of the group parietal mask **(Supplementary, Figure S4)**. The TMS coil was then oriented around the scalp normal vector so that the direction of the second phase of the induced E-field coincides with the inward-pointing normal vector on the sulcal wall closest to the brain target location. This pulse direction induces the strongest E-field and activation at the target (Kammer et al., 2001; Janssen, Oostendorp, & Stegeman, 2015). The sulcal wall was identified using Freesurfer’s gyral/sulcal cortex classification (Fischl et al., 2004: file lh.sulc, a byproduct of SimNIBS’ mri2mesh script during the brain surface extraction) and a brain surface point was chosen at the transition location in-between local concavity and convexity (|local curvature threshold| < 0.05) defining the sulcal wall. In order to compute the normal vector of that sulcal wall point, the surface normal of the triangles were averaged. The intended coil orientation was then entered in the neuronavigation system using the ‘twist’ tool.

### 2.5. Stimulation amplitude approach

Rather than defining rTMS pulse amplitude according to a percentage of the motor threshold, as is frequently done in the literature, amplitude here was defined according to target-specific E-field values. While the motor threshold provides individualized information regarding the cortical reactivity of the motor cortex, it does not take into account differences in head anatomy and brain physiology between motor cortex and other cortical regions within an individual. As such, traditional amplitude calibration based on the motor threshold may lead to substantial variation in the desired E-field strength in the targeted brain region. This may lead to response variability, since the E-field strength is the key determinant of neural activation by TMS (Aberra, Wang. Grill, & Peterchev, 2018). Therefore, in the present study we selected the TMS pulse amplitude (coil current rate of change, di/dt) to induce a specific E-field magnitude, E_ref_, in the left lateral parietal region of interest (ROI) across subjects.

To define E_ref_, computer simulations were used to estimate the E-field distribution within the parietal ROI induced when TMS was applied at amplitude equal to the resting motor threshold in each of 9 subjects from a previous study (Beynel et al., 2019, **Supplementary, Figure S2**). For each of the 9 subjects, a parietal ROI was constructed by individual fMRI activity (|z|>0; within a group activity mask, Beynel et al., 2019) registered to the individual’s space (FSL *flirt*, Smith et al., 2004) within voxels classified as gray matter (SimNIBS: gm_only.nii.gz). The selected voxel were ranked according to their E-field strength, and a metric, E_100_, was defined as the minimum strength across the 100 voxels with strongest E-field (**Supplementary, Figure S3**). The meaning of this metric is that 100 voxels in the ROI, corresponding to a volume of 100 mm^3^, have E-field strength larger than E_100_. The average E_100_ across the 9 subjects was calculated to be 56 V/m, which we set as our desired target E-field strength, E_ref_ = 56 V/m.

To select the individual TMS pulse amplitude in this study, computer simulations were performed to estimate the individual E-field distribution (analyzed for the left parietal ROI) and determine a TMS coil di/dt for which E_100_ = E_ref_ in the ROI for each subject. Since TMS pulse di/dt scales linearly with the induced E-field, TMS was simulated for di/dt = 10^6^ A/s, and a scaling factor was computed for di/dt to reach E_ref_ for the hair thickness measured during the screening visit. The individual’s di/dt-value was determined for different hair thicknesses (in steps of 0.5 mm from scalp surface) and stored in a reference table (**Supplementary Table 1**). During the first TMS visit, the hair thickness at the exact stimulation location was re-measured and rounded to match the closest value in the table. The corresponding computed di/dt value was selected. The TMS amplitude, expressed as a percentage of the maximum stimulator output (MSO) was adjusted for the chosen di/dt value. The amplitude, together with the determined location and orientation (described in section 2.4) were then experimentally applied. Two E-field strengths in the targeted region were experimentally tested, with E_100_ metric equal to either 80% E_ref_ or 100% E_ref_. Resting motor threshold assessed during the screening visit was used to ensure that all stimulation intensities were below 130% of the resting motor threshold, and therefore within the published safety guidelines (Rossi, 2009).

The computer simulations of the TMS-induced E-field were performed using the SimNIBS software package (Version 2.0.1, Thielscher et al., 2015). A computational model of each participant’s head was first generated employing co-registered T1- and T2-weighted MRI data sets to model major head tissues (scalp, skull, cerebrospinal fluid, gray and white brain matter) represented as tetrahedral mesh elements. Each mesh element was assigned a conductivity value based on its tissue association. The scalp, skull, and cerebrospinal fluid conductivities were set to isotropic values of 0.465, 0.010, and 1.654 S/m, respectively. The gray and white matter compartments were assigned anisotropic conductivities to accounting for the fibered tissue structures. This was accomplished within SimNIBS by co-registering diffusion weighted imaging data (available for 14 out of 15 participants) employing a direct mapping approach (e.g., Rullmann et. al, 2009) to establish the required conductivity tensors and scale their magnitude up to default literature-based values of 0.275 and 0.126 S/m for gray and white matter, respectively. For the three subjects with missing DTI information, the latter values were assigned as isotropic conductivities. (**Supplementary Table 2** for individual subjects ‘information)

### 2.6. TMS procedure

During visits 3 to 6, participants performed the DRAT while active or sham rTMS was delivered to the individualized left lateral parietal target using an A/P Cool-B65 coil (MagVenture, Alpharetta, GA, USA). Twenty-five pulses of 5 Hz rTMS were delivered during the delay period of each trial (Figure 1). For every two trials with stimulation, one trial without stimulation was performed. This approach, successfully used in multiple studies by Luber et al. (2007, 2008, 2013), theoretically allows time for neural activity in the stimulated region to return to its homeostatic baseline, allowing for greater range for production of rTMS-induced plasticity and thus, potentially, greater rTMS effect on behavioral performance. The non-stimulated trials were excluded from subsequent analyses. The two intensities of stimulation, 80% E_ref_ and 100% E_ref_, were applied on different days. Sham stimulation was applied using the same coil in placebo mode, which produced clicking sounds and somatosensory sensation via electrical stimulation with scalp electrodes similar to the active mode, but without a significant E-field induced in the brain (Smith, & Peterchev, 2018). This type of sham stimulation allows participants to stay blinded during the course of the experiment. Neuronavigation (BrainSight, Rogue Research, Canada) and real time robotic control (SmartMove, ANT, Netherlands) were used to ensure that the optimal coil position was maintained throughout the stimulation sessions.

On each TMS visit, subjects performed the DRAT at their two individually titrated difficulty levels (Easy and Hard). Ten blocks of the DRAT task were performed (30 trials per block): one block without stimulation (No-Stim1), followed by five blocks of active or sham stimulation, one block without stimulation (No-Stim2), and five more blocks with the sham or active stimulation. The first 5 rTMS blocks in the first visit were always active rTMS at 100% E_ref_ to ensure that the subjects were able to tolerate this stimulation amplitude, with the later 5 rTMS blocks being sham stimulation at output setting corresponding to the 100% E_ref_ condition. For the three other visits, the intensities of stimulation were alternated by day and sham and active stimulation were applied on the same day in a random order. No adverse events or pain due to the stimulation were reported by any of the subjects. As noted above, our central hypothesis was that older adults would show a benefit for WM accuracy on the DRAT due to online rTMS, but only during the most difficulty condition.

### 2.7. Statistical Analyses

Analyses were performed using the general linear model module of Statistica (TIBCO Software, Palo Alto, USA), normality was tested using Kolmogorov-Smirnoff tests, and multiple comparisons corrections were performed using Bonferroni correction. All the results are expressed as mean ± standard error.

## 3. Results

### 3.1. Performance without rTMS

Four individualized difficulty levels (Very Easy, Easy, Medium, and Hard) were defined according to participant’s performance on the staircase version of the DRAT performed during Visit 1. At the Very Easy level, the absolute number of letters to maintain and alphabetize was found to be: 3 (n=11), 4 (n=3) or 5 (n=1). During the rTMS sessions on Visits 3 through 6, only Easy and Hard difficulty levels were used. To ensure that these difficulty levels were properly defined, to test the differences between Valid and Invalid trials, and to assess learning across time, repeated measure ANOVA was performed with the following within-subject factors: Visit (Visit 3, Visit 4, Visit 5 and Visit 6), Difficulty (Easy and Hard) and Task Condition (Valid and Invalid trials). To prevent contamination of the data due to potential rTMS carryover effects, only behavior obtained during the first block without stimulation (No-Stim1) was considered. Trials for which the subjects did not answer (1.79 ± 0.57%) were excluded. A significant main effect of Difficulty was found (F(1, 13)=112.55, p<.001) with higher accuracy for Easy (88.86 ± 3.32%) than for Hard (63.43 ± 3.25%) difficulty levels. No significant main effect of Visit F(3, 39)=2.17, p=.11), Task Condition (F(1,13)<1) or interaction between these factors were found. As such, it can be inferred that the difficulty levels were well defined by the staircase procedure. Subjects did not exhibit significant learning across visits, and accuracy was equivalent in the Valid and the Invalid trials.

### 3.2. TMS dosing results

As described in the methods section, TMS targeting used individualized coil position, coil orientation, and stimulation amplitude based on fMRI, structural MRI, and E-field data. The following sections will therefore present the individual and group results for each of these parameters.

#### 3.2.1 TMS coil position and orientation

TMS coil position was constrained by a group mask defined on the left lateral parietal cortex (MNI coordinates at center of mask: −41; −64; 42, Figure 3A). For each subject, the peak fMRI activation associated with difficulty increased during the DRAT within this mask and was used to define the coil position. TMS coil orientation was then selected such that the second phase of the induced E-field was perpendicular and directed into the nearest sulcal wall for each individual subject. Figure 3B illustrates the final coil position and orientation for each subject.

**Figure 3:**
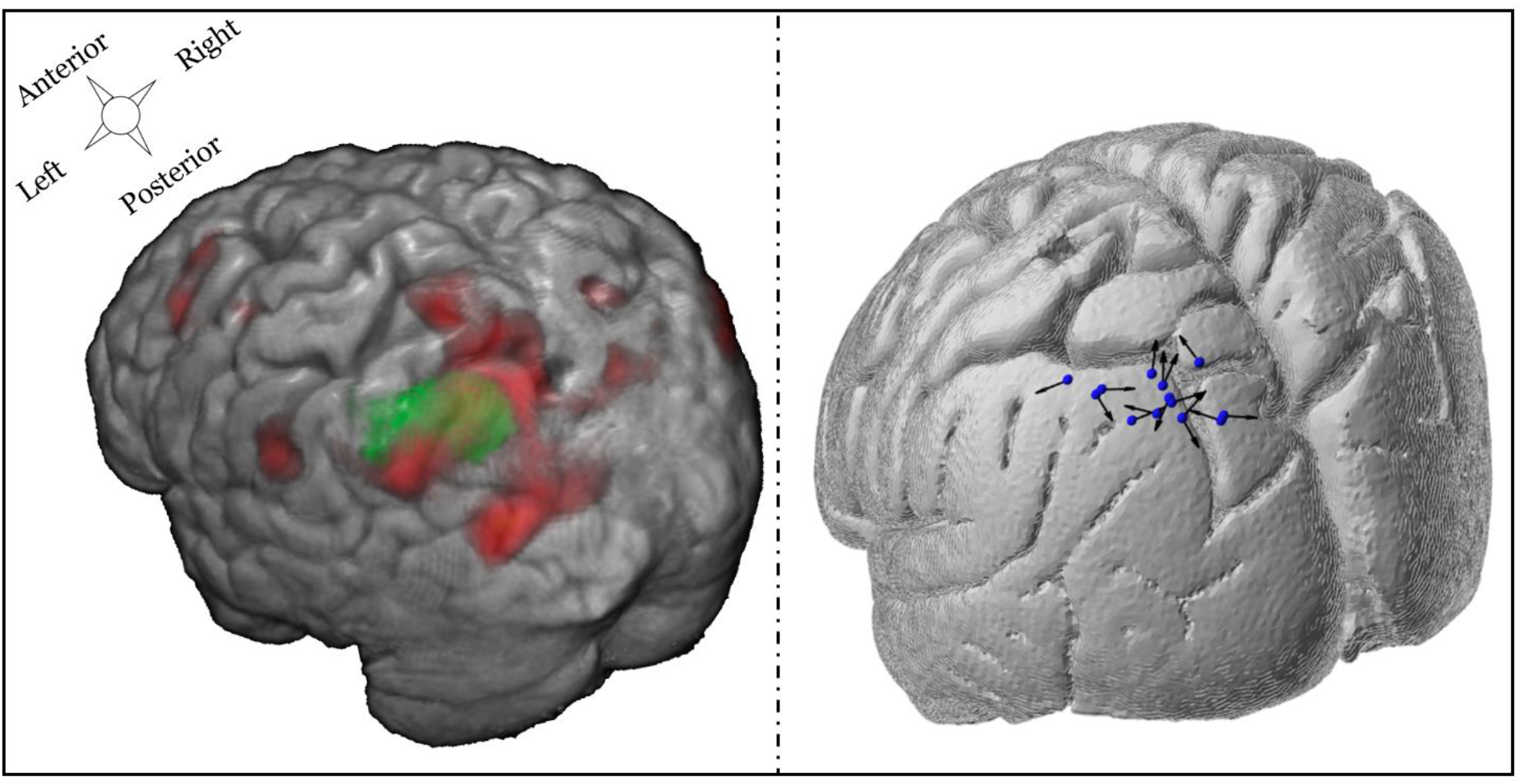
Left panel: Example of one individual subject combining the group mask over the left parietal cortex (green) combined with individual fMRI activation (red) associated with the parametric increase in difficulty during the delay. Right panel: TMS coil position and orientation for each participant. The blue spheres represent the coil location and the black arrows correspond to the direction of the second phase of the induced E-field pulse (some of the arrowheads are not visible because of the 3D view).

#### 3.2.2 Stimulation Amplitude

Figure 4 shows the relationship between the individual TMS pulse amplitude inducing 100% E_ref_ at the parietal target and the resting motor threshold. This correlation was positive but not significant (r = 0.38, p = 0.16). The positive slope of the regression line likely reflects the fact that the motor threshold captures some anatomical factors such as scalp-to-cortex distance that are correlated, to some degree, between various brain regions—this is the premise of conventional motor threshold-based dosing. The lack of significance in the correlation, however, supports the premise of our dosing approach, specifically that matching the E-field exposure of the target across subjects results in pulse intensities that are not predicted by motor cortex reactivity. The pulse amplitude based on E-field modeling was below resting motor threshold for some subjects (n=8), while it was above this threshold for the others (n=7). This inter-subject variability suggests that studies which dose based on a percentage of resting motor threshold may result in considerable within-group variability in the actual E-field that is induced in the cortex. In the current study, this potentially unwanted variability is controlled through appropriate dosing that considers the magnitude of the E-field in the desired cortical target. It should be noted that the resultant individual pulse intensities did not exceed the rTMS safety guidelines of < 130% of resting motor threshold for trains of less than 10 seconds in any of the subjects (Rossi et al., 2009).

**Figure 4:**
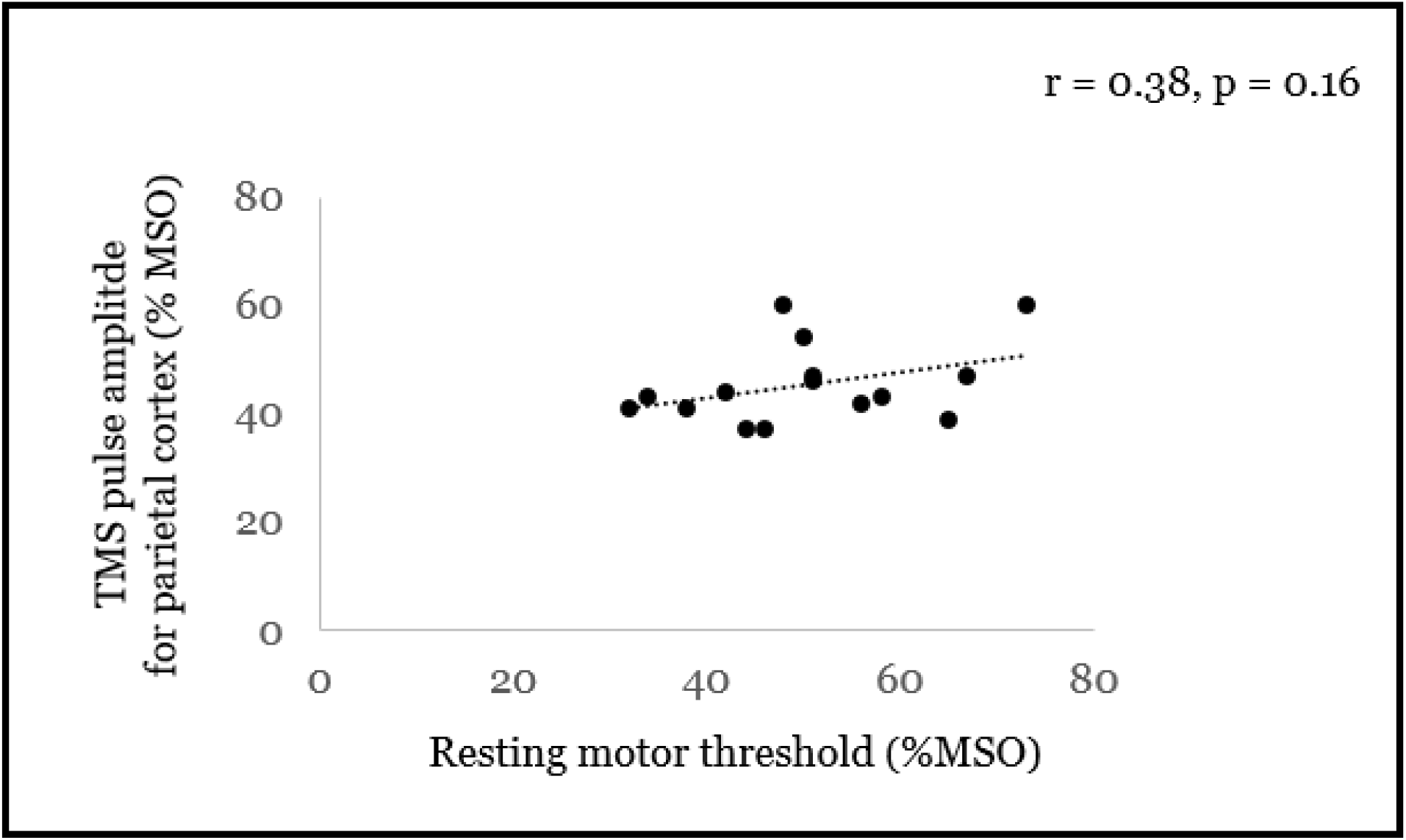
Relationship between the TMS pulse amplitude determined by modeling to induce uniform E-field exposure of parietal target across subjects, and the resting motor threshold. Both are expressed as a percentage of the maximum stimulator output, MSO. No significant correlation was found.

### 3.3. rTMS effects

#### 3.3.1. Cumulative rTMS effects

In the current design, active and sham stimulation were performed on the same day. To ensure that no cumulative carryover effects contaminated the effect of one type of stimulation over the other, the block of trials performed either immediately before (No-Stim1) or immediately after each type of stimulation (No-Stim2_AfterActive, and No-Stim2_AfterSham) were analyzed separately. Trials for which the subject did not respond were excluded (1.70 ± 0.49 %). One way ANOVA, performed on accuracy across these three blocks, did not reveal significant differences between blocks performed before stimulation (No-Stim1= 76.38 ± 2.36 %), blocks performed after active rTMS (No-Stim2_AfterActive= 76.29 ± 2.05 %), or blocks performed after sham rTMS (No-Stim2_AfterSham= 78.08 ± 2.32; F(2,28)<1). This result suggests that no carryover effects persisted and thus subsequent rTMS effects were not contaminated by the former type of stimulation.

#### 3.3.2. Omnibus rTMS effects on the DRAT

To assess rTMS effects on performance during the DRAT, repeated measures ANOVA was run on accuracy scores with the following within-participant factors: Task Condition (Valid and Invalid), Stimulation Type (Active and Sham), Stimulation Amplitude (80 % and 100 % E_ref_), and Task Difficulty (Easy and Hard). Trials for which the subjects did not answer (1.93 ± 0.4%), and trials during which stimulation was not applied (33.3 % No-Stim trials), were excluded from this analysis.

This ANOVA produced non-significant main effects of Task Condition (Valid: 72.84 ± 6.53 %, Invalid: 71.24 ± 5.86 %, F(1,14) < 1) and Stimulation Type (Active: 71.70 ± 6.29 %, Sham: 72.38 ± 6.12 %, F(1,14)<1) on task accuracy. A significant main effect of Stimulation Amplitude was observed, however, with lower accuracy when stimulation was applied at 100% E_ref_ (70.19 ± 6.26 %) compared to stimulation applied at 80% E_ref_ strength (73.89 ± 6.12 %; F(1,14) = 10.15, p = 0.007, η^2^ = 0.42). A significant main effect of Difficulty was also observed (F(1,14) = 117.43, p <0.001, η^2^ = 0.89), with lower accuracy for hard trials (56.59 ± 5.60 %) compared to easy trials (87.49 ± 3.6 %). A significant three-way interaction was also found between Task Condition, Stimulation Type and Stimulation Amplitude (F(1,14) = 5.14, p = 0.04, η^2^ = 0.27). The decomposition of this interaction revealed that, only for the Invalid trials, active rTMS applied at 100% E_ref_ (69.14 ± 6.4 %) tended to disrupt accuracy compared to active rTMS applied 80% E_ref_ (74.25 ± 6.1 %) (F(1,14) = 4.19, p = 0.088). This suggests that applying rTMS at a stronger amplitude leads to a larger rTMS effect. ANOVA also revealed a significant interaction between Stimulation Type and Difficulty (F(1,14) = 9.70, p = 0.008, η^2^ = 0.41). Bonferroni corrected post-hoc comparisons revealed that while no differences were found between active (87.97 ± 3.7 %) and sham stimulation (87.00 ± 3.71 %) for Easy trials (F(1,14) = 2.22, p = 1), for Hard trials active rTMS significantly decreased accuracy (55.42 ± 5.50%) compared to sham rTMS (57.76 ± 5.70 %; F(1,14) = 4.12 p = 0.045; Table 1). This result indicates that rTMS only disrupts performance for harder memory trials.

**Table 1:** Average accuracy for the Difficulty levels and the Stimulation Type.

|  | Easy | Hard |
| --- | --- | --- |
| Active rTMS | 87.97 ± 3.70 % | 55.42 ± 5.50 % |
| Sham rTMS | 87.00 ± 3.71 % | 57.76 ± 5.70 % |
| p-values | NS | 0.045 |

#### 3.3.3 Follow-up tests on the effects of stimulation amplitude

To further investigate differences between stimulation applied at 80% and 100% E_ref_, a separate follow up ANOVA was performed with the factors Difficulty (Easy and Hard) and Stimulation Type (Active and Sham) for each stimulation amplitude. Results for 100% E_ref_ showed that accuracy was significantly decreased in Hard trials (60.02 ± 2.65 %), relative to Easy trials (88.38 ± 2.65 %; (F(1,14)=148.33, p <0.001, η^2^ = 0.91). While the main effect of Stimulation Type was not significant (Active: 73.37 ± 4.72 % vs. Sham: 75.04 ± 4.29 %, F(1,14) = 2.89, p = 0.11, η^2^ = 0.17), there was a significant interaction between Stimulation Type and Difficulty (F(1,14) = 6.42, p = 0.02). Post-hoc Bonferroni comparisons showed that, while no differences were found between active and sham stimulation at the Easy difficulty level (Active: 88.49 ± 2.79 % vs. Sham: 88.28 ± 2.61 %, p = 1), active rTMS did significantly decrease accuracy (58.26 ± 2.39 %) compared to sham rTMS (61.79 ± 2.40 %; (F(1,14) = 6.68, p = 0.03) on the Hard trials.

For the ANOVA performed with rTMS applied at 80% E_ref_, a main effect of Difficulty was found with lower accuracy for Hard trials (63.54 ± 2.28 %) compared to the Easy ones trials (90.84 ± 2.24 %, F(1,14) = 136.0, p < 0.01, η^2^ = 0.91). No differences were found for Stimulation Type (F(1,14)<1). Interestingly, the interaction between Difficulty and Stimulation Type was not significant for this stimulation amplitude (F(1,14) = 2.02, p = 0.17). These results therefore show that the disruptive effects of rTMS for harder memory trials are specific to the highest stimulation amplitude. However, this effect needs to be interpreted with caution because, as indicated by the non-significant interaction between E-field Strength, Stimulation Type and Difficulty (F(1,14)<1), in the larger ANOVA, it did not reach omnibus significance.

At the individual level, when computing the rTMS effect as a percentage of change between active and sham rTMS the results showed that when stimulation was applied at 100% E_ref_, 9 out of 15 subject showed performance disruption, while 7 showed disruption when rTMS was applied at 80% E_ref_ (Figure 5).

**Figure 5:**
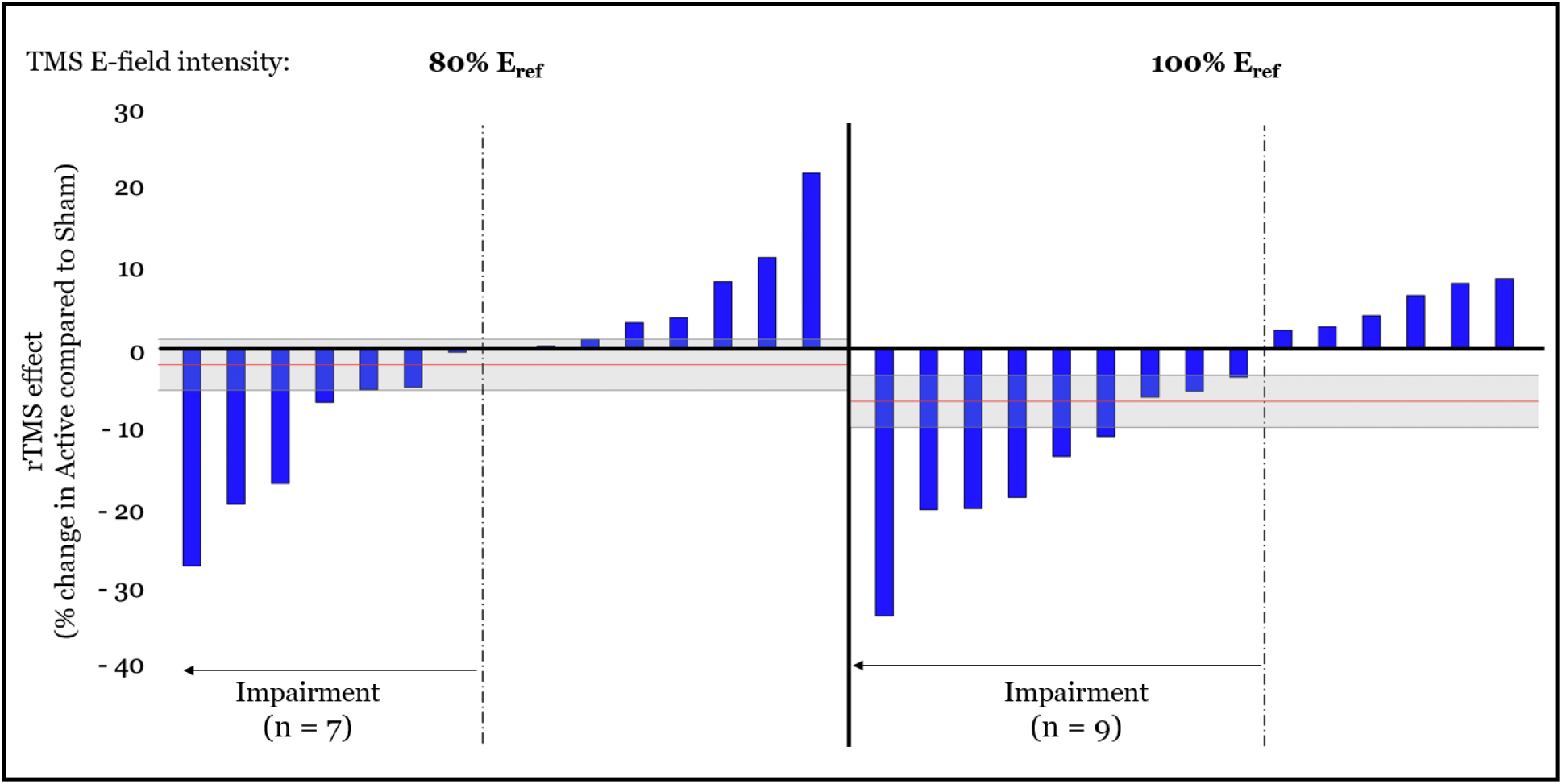
rTMS effect (expressed as a percentage of change between Active and Sham rTMS) for rTMS applied at 80% E_ref_ (left) and at 100% E_ref_ (right). Each blue bar represents one subject. The red horizontal line shows the averaged rTMS effect across each subject (mean= −2.03 %, and −6.66 % for 80% E_ref_ and 100% E_ref_, respectively). The gray rectangle represents the standard error (SE = 3.19 % and 3.26 % for 80% E_ref_ and 100% E_ref_, respectively).

### 3.4. Effect sizes comparison with prior study

As mentioned in the introduction, the goal of the current study was to optimize the faciliatory effects of rTMS observed in Beynel et al 2019, where 5Hz rTMS was applied over the left dorsolateral prefrontal cortex. Here, we wished to maximize the difference between active and sham rTMS by stimulating the parietal cortex, which has been shown to be more activated during the DRAT (Davis et al., 2018), and by modifying some stimulation parameters. To compare the effect sizes for these two studies, Cohen’s d was calculated from the individual conditions that yielded the largest effects in each study (hardest difficulty level in the Invalid trials in Beynel et al. (2019) and 100% E_ref_ strength for the current study). Cohen’s d was calculated as follows:

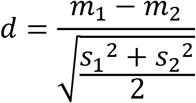

where *m*_1_ and *m*_2_, and *s*_1_ and *s*_2_ are the mean and standard deviation of accuracy for active and sham, respectively.

Results show a modest rTMS-induced effect size for both studies and highlight the opposite directionality of this effect. While the effect size was negative (Cohen’s d = −0.38) in Beynel et al. (2019), it is positive in the current study (Cohen’s d = 0.34). This contrast illustrates that, instead of optimizing the facilitatory rTMS effect obtained in the former study, modifications made in the current study reversed the rTMS effect, while producing relatively equivalent effect sizes.

## 4. Discussion

This study was conducted to test whether parameter-optimized rTMS, delivered to the left lateral parietal cortex, could enhance working memory manipulation in healthy elderly subjects. In our previous study (Beynel et al., 2019), applying 5Hz rTMS over the left prefrontal cortex increased young and elderly participants’ accuracy, but the effect was small. Therefore, in the current study, the goal was to modify the stimulation target and optimize stimulation parameters in order to produce larger performance enhancements. However, these changes led to an opposite pattern of findings wherein active stimulation yielded a small performance impairment relative to sham stimulation, with a similar effect size. The following sections discuss these patterns of effects, as well as the innovative targeting and dosing approaches used in this study.

### 4.1. Site- and timing-specific rTMS effects on WM manipulation

In our studies, online rTMS was applied with a goal of enhancing the manipulation of information in WM, as assessed by performance on the DRAT. As reported in Davis et al. (2018), group analysis of fMRI acquired during the second visit of these studies revealed that when participants are mentally maintaining and alphabetizing letters during the delay period of the DRAT, the left lateral parietal cortex produced strong activation. This finding is consistent with a role of parietal cortex in symbolic computations (Piazza et al., 2007; Dehaene et al., 2003; Park et al., 2013) and led to targeting of the parietal cortex, rather than DLPFC, in the current study. As such, contrasting findings between the two cohorts reflect site specificity of rTMS effects on WM.

While it is widely reported in the literature that online rTMS applied during a task induces a temporary “virtual lesion”, evidence of performance enhancement has also been found (see Luber & Lisanby, 2014 for a review). One factor likely mediating the opposite effects of online rTMS is the modulation of endogenous task-related oscillatory dynamics in a manner that is specific to the timing of ongoing processes (Johnson, Hamidi, & Postle, 2010). Working memory tasks, in particular, have been associated with 5Hz theta-band coupling between frontal and parietal regions during the memory retention period, which increases parametrically with memory load (Jensen, & Tesche, 2002). However, contrary to such expectations, applying 5Hz rTMS during the delay period *disrupted* participants’ accuracy and suggest a more complex interaction between site-specific and timing-specific rTMS effects. One possible explanation for this discrepancy is that while rTMS may engender certain oscillatory patterns at the site of stimulation, there exists some general latency after the end of the last TMS pulse for that entrainment to emerge. Thus, rTMS applied to a region during the time it is engaging in task-essential processing will disrupt performance, while stimulation trains of appropriate frequency applied prior to that processing can enhance performance, possibly due to entrainment of functional oscillations prior to the essential cortical activity (Hanslmayr et al., 2014; Roberts et al., 2018). The fMRI evidence suggests the essential processing in parietal cortex needed for the DRAT occurred during the delay period, with rTMS applied in that period injecting random noise that disturbed performance, while delay period 5 Hz rTMS to DLPFC may result in pre-processing activity-possibly through entrainment of theta frequency-that may enhance processing there during the subsequent probe period. As such, applying rTMS over the parietal cortex, before the encoding might lead to performance enhancement; however, more investigation is needed to determine the stimulation timing parameters that will lead to such enhancement.

### 4.2. Electric-field-based TMS dosing

We adopted a novel individualized TMS targeting strategy that combined fMRI, structural MRI, and E-field modeling to select the TMS coil position, orientation, and pulse amplitude. The TMS coil was centered over the individualized peak fMRI activation, within a group mask based on previously reported fMRI data for this task (Beynel et al., 2019), and associated with difficulty level in the DRAT. The subject’s brain anatomy, imaged via MRI, was then used to define the optimal coil orientation such that the E-field pulse was perpendicular and directed into the nearest sulcal wall. As such, this approach allowed us to address the potential confounds associated with cortical thinning associated with older adult samples, and builds on previous research in the motor cortex showing lowest threshold for motor evoked potentials when the induced E-field is perpendicular and flowing into the sulcal wall (Brasil-Neto et al., 1992; Kammer et al., 2001; Richter et al., 2013). This observation has been replicated in various other cortical regions (Hill, Davey, & Kennard, 2000; Janssen, Oostendorp, & Stegeman, 2015; Kammer et al., 2001). The likely explanation for the optimality of current perpendicular to the sulcal wall is that this current orientation induces the strongest E-field in the corresponding gyrus (Janssen, Oostendorp, & Stegeman, 2015). The likely reason for the sensitivity to current direction (current flowing into the gyrus) is the morphology and orientation of pyramidal neurons in the cortex (Aberra et al., 2018). Furthermore, we found no effect of cortical thickness across subjects on the observed TMS-related effects on performance, suggesting that E-field modeling is a good method to prevent this bias (**Supplementary, Figure S5**). Since these observations appear to be generalizable to any area of cerebral cortex, we adopted this approach in our study as a means to minimize the pulse amplitude required for stimulating the target and to maximize the stimulation focality.

The traditional approach for individualizing the amplitude of rTMS pulses is to scale the coil current based on the easily-observable resting motor threshold response. While this approach is appropriate for stimulation of primary motor cortex, where it captures the underlying anatomical and physiological variability across subjects, its use in other brain regions has significant limitations. First, the individual location of the muscle representation in primary motor cortex affects the motor threshold but is unlikely to match the location of the non-motor target area on its respective gyrus. Second, the local anatomy of the head in the vicinity of the target can vary individually. Third, the local physiology may differ from that of motor cortex in an individual manner. Therefore, we opted to match the delivered E-field strength at the cortical target across subjects. The reference E-field strength, E_ref_, was selected based on estimates of the average stimulation delivered in prior studies with a similar target. We introduced a metric, E_100_, that ensures that a fixed volume of the cortical target is exposed to an E-field strength at or above the reference value, while also reducing the sensitivity to computational outliers that may impact other E-field metrics such as peak or mean value. While the design of the current study did not compare the effect of rTMS applied with dosing based on the motor threshold versus the simulated E-field strength, the results highlighted a significant difference between stimulation applied at 80% or 100% of E_ref_. Indeed, stronger disruptive rTMS effects were associated with stimulation applied at higher amplitude. This suggests that the E-field-based dosing approach produced sufficiently consistent rTMS effects across subjects to differentiate the two amplitude conditions. More investigation is needed to explore the value of this new dosing approach.

## 5. Conclusion

5 Hz rTMS to left lateral parietal cortex during the delay period of the DRAT was found to impair WM behavior of healthy older adults. When considered in light of Beynel et al 2019, the effects can be interpreted as site- and timing-specific effects of rTMS on WM manipulation. Collectively, across these two cohorts, findings demonstrate the ability of online rTMS to up-regulate and down-regulate WM by stimulating prefrontal and lateral parietal cortices, respectively. This information is important for further clarifying how rTMS may be used to therapeutically treat disorders where memory is impacted. Future investigation is warranted to refine the parameters under which these effects are present, and to explore how targeting and dosing considerations can be adjusted to optimize rTMS efficacy.

## 6. Funding

This research was funded by grant U01 AG050618 from the National Institute of Aging (https://www.nia.nih.gov/) and in part by the Intramural Research Program of the National Institute of Mental Health (https://www.nimh.nih.gov/index.shtml).

## Declaration of interest

A. V. Peterchev is inventor on patents and patent applications and, in the past 5 years, has received travel support as well as patent royalties from Rogue Research; research and travel support, consulting fees, as well as equipment donation from Tal Medical / Neurex; research and patent application support from Magstim; as well as equipment loans from MagVenture, all related to TMS technology, but not directly related to the presented work.

