## supplementary material for "Site-specific effects of online rTMS during a working memory task in healthy older adults"

### TMS dosing method

In this study, the stimulation target was defined by combining individualized fMRI activation data to define the position of the coil center, anatomical MRI data to define the coil orientation and pulse direction, and E-field modeling to define the TMS pulse amplitude, which to our knowledge has never been implemented before. These steps are summarized below.

#### Coil position

The position of the coil center was defined to be on the scalp directly over the individual cortical target. The cortical target was defined as the peak activation within the left parietal cortex associated with the increase in difficulty during the delay period of the delayed-response alphabetization task (DRAT). To constrain the stimulation target, we used a parietal group mask based on activation in 22 older adults who participated in our previous study (Beynel et al., 2019) (see Figures 1 and 3A in the paper).

#### Coil orientation

The TMS coil orientation was defined such that the coil handle, which is parallel to the primary induced E-field under the center of the coil, was perpendicular to the sulcal wall nearest to the cortical target. The sulcal wall was automatically determined using the FreeSurfer software and the inflection point of the wall curvature closest to the cortical target was identified. The normal vector of that sulcal wall point was defined by averaging the surface normals of surrounding surface mesh triangles and used to orient the coil so that the second phase of the induced E-field pulse was directed into the gyrus (see Figure S1 and Figure 3B in the paper).

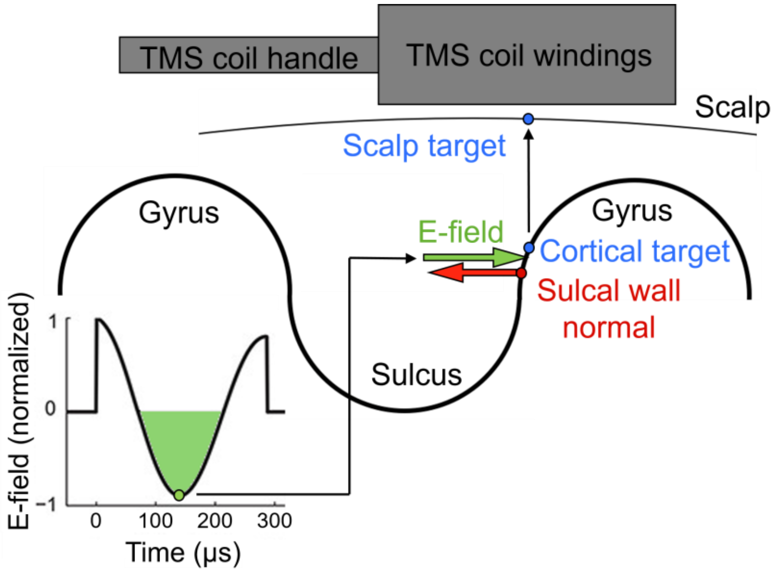


**Figure S1**. Targeting approach to locate and orient the TMS coil. Based on the individual cortical target (depicted in green), the closest locations on the scalp (depicted in blue) as well as at the inflection point on the gyral wall (depicted in red) are determined. The surface normal vector at the gyral wall point is used to orient the (longitudinal axis) of the coil so that their angle equals to 180 degrees. In that way, the second phase of the induced E-field is directed into the gyral wall.

### TMS pulse amplitude

The TMS pulse amplitude was defined according to site-specific E-field values. This process allows standardizing the strength of induced E-field entering the cortical target across all subjects. Before the study starts, a reference E-field strength ($E_{ref}$) was defined. To do so, computer stimulations were used to estimate the E-field distribution within the parietal mask (Figure S2) that was induced when TMS was applied at 100% rMT for 9 subjects from our prior study (Beynel et al., 2019). The E-field value of the voxel ranked 100 by E-field strength ($E_{100}$) was determined within the parietal group mask for the 9 subjects (see Figure S3). The target E-field strength used in the current study was then computed as the averaged E100 across these 9 subjects ($E_{ref}$ = 56 V/m).

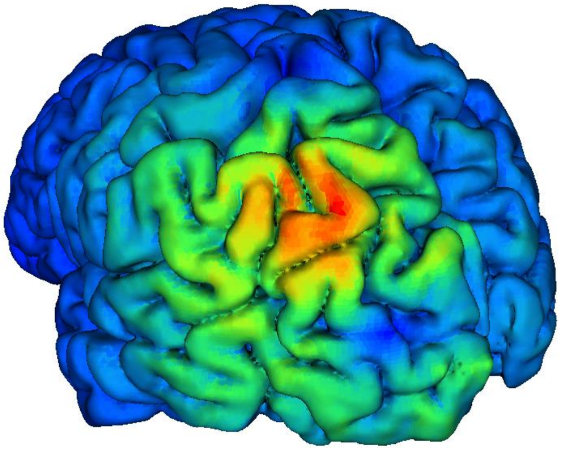

**Figure S2**. Example of the E-field magnitude plotted on the cortical surface, when targeting the left parietal ROI.

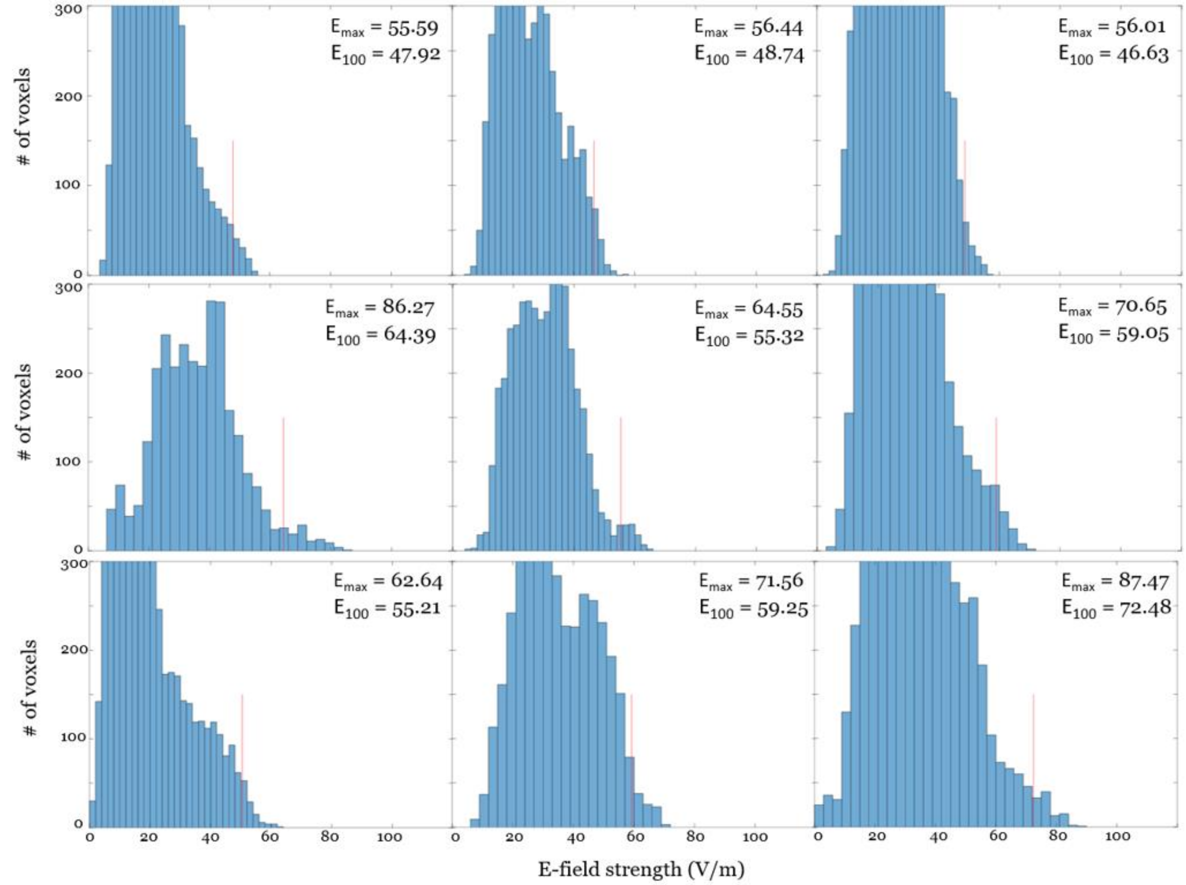


**Figure S3.** TMS E-field strength histograms for each of 9 subjects from our prior study (Beynel et al. 2019) and their corresponding $E_{100}$ value when targeting parietal cortex at 100% of the individual resting motor threshold. These $E_{100}$ values were used to compute $E_{ref}$ in the current study.

For each subject in the current study, computer simulations were performed to estimate the individual E-field distribution and determine the appropriate TMS pulse amplitude (di/dt) to reach $E_{100}$ of 56 V/m. The individual di/dt value was then determined for different hair thicknesses (in steps of 0.5 mm from the scalp surface) and stored in a reference table (see Table S1 for an example).

| Hair thickness (mm) | 0 | 0.5 | 1 | 1.5 | 2 | 2.5 | 3 | 3.5 |
|---|---|---|---|---|---|---|---|---|
| di/dt (A/µs) | 70.2 | 71.9 | 73.6 | 75.4 | 77.2 | 79.0 | 80.9 | 82.8 |

**Table S1**. Example of a reference table showing the di/dt values for different hair thicknesses for one subject (ID: S237).

When the subject arrived for the first TMS visit, compressed hair thickness was measured by placing the BrainSight pointer over the individualized TMS target on the subjects' head, and positioning the depth gauge mounted on a plastic plate at this location (Figure S4). As shown in Table 2, hair thickness is an important factor to consider given the large variability between subjects (from 0 to 8 mm). The closest hair thickness value in the reference table was found, and the corresponding di/dt value was chosen. Finally, the stimulator pulse amplitude (%MSO) was then adjusted so that the di/dt reported by the stimulator matched the desired value. The subject-specific stimulation information is provided in Table S2.

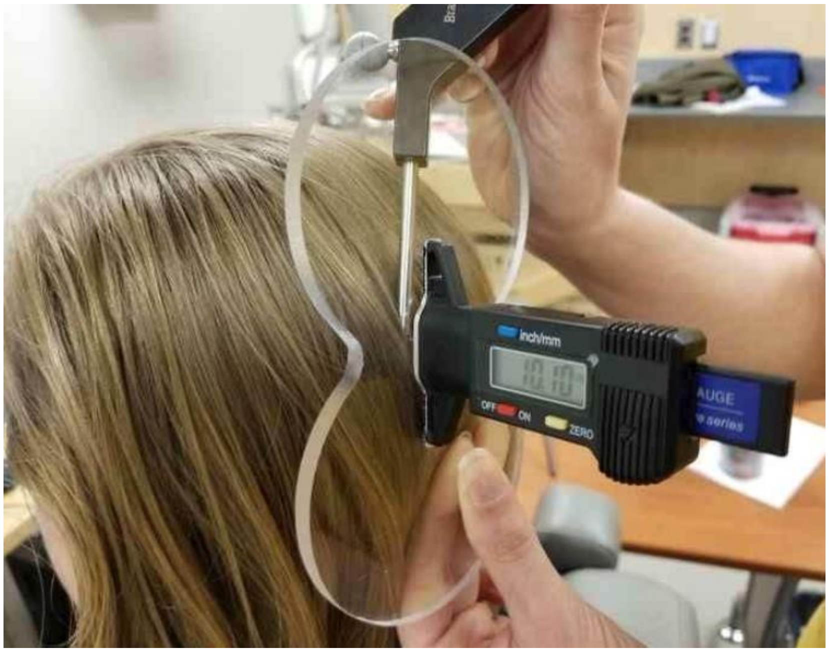


**Figure S4**. Illustration of the process of measurement of compressed hair thickness.

| Subj. ID | rMT | | TMS coil placement | | Hair thick-ness (mm) | Pulse amplitude | |
|---|---|---|---|---|---|---|---|
| | %MSO | di/dt (A/µs) | Position (mm) | Orientation (°) | | % MSO | di/dt (A/µs) |
| 236 | 58 | 87 | −42.6, −24.4, 0.98 | −110 | 2 | 43 | 66.4 |
| 237* | 67 | 106 | −40.9, −17.5, 14.5 | 56.2 | 0.5 | 47 | 71.9 |
| 240 | 32 | 48 | −26.3, −37.6, −8.26 | 62.3 | 1 | 41 | 62.6 |
| 244 | 73 | 111 | −35.7, −18.2, 35.2 | −129 | 8 | 60 | 89.8 |
| 247 | 34 | 48 | −29.4, −37.3, 9.60 | −166 | 1 | 43 | 64.7 |
| 248 | 44 | 66 | −34.2, −39.0, −15.2 | 157 | 1 | 37 | 54.6 |
| 249 | 48 | 73 | −25.9, −38.4, 7.13 | −130 | 1.6 | 60 | 90.4 |
| 250* | 38 | 56 | −32.3, −33.2, 15.3 | 150 | 2 | 41 | 61.9 |
| 251 | 42 | 62 | −37.3, −29.5, 12.2 | −45.4 | 2.2 | 44 | 66.1 |
| 252 | 46 | 69 | −30.6, −45.9, −19.7 | 10.4 | 0 | 37 | 56.8 |
| 262 | 51 | 76 | −25.2, −41.8, 11.4 | 118 | 0 | 46 | 70.2 |
| 263 | 50 | 75 | −53.8, −11.5, 14.2 | 1.31 | 0 | 54 | 81.8 |
| 264* | 56 | 84 | −32.8, −29.2, 7.20 | 133 | 1.5 | 42 | 63.4 |
| 266 | 51 | 76 | −27.0, −33.6, 7.68 | 83.3 | 1 | 47 | 70.8 |
| 269 | 65 | 101 | −37.1, −27.3, −7.16 | 83.8 | 3 | 39 | 57.2 |

**Table S2**. Subject ID, resting motor threshold (rMT), TMS coil placement parameters, hair thickness at the target, and pulse amplitude for the 100% $E_{ref}$ rTMS condition. The TMS coil position is the location of the center of the coil in the individual space, and the coil orientation is relative to the midline. The asterisk next to the subject IDs indicates that no DTI was acquired. The discrepancy between the MSO and the di/dt for example between subjects S236 for which 43% MSO corresponds to 66.4 A/µs, and S241 for which 44% MSO corresponds to 66.1 A/µs is due to some variability in the TMS device di/dt reading.

### Impact of cortical thickness on rTMS effect

As aging has been shown to be associated with structural changes such as steady linear decline in cortical thickness, we performed an exploratory correlation between the TMS effect obtained in the hardest difficulty level of the DRAT and the subject's cortical thickness at the cortical target. The stimulated region was mapped on FreeSurfer, and cortical thickness was extracted from this ROI. No correlation was found between these two factors (Figure 5). As can be seen in this figure, there was an outlier with an rTMS effect more than two standard deviations away from the mean. After removing this data point, the correlation was still not significant ($r = -0.50$, $p = 0.07$).

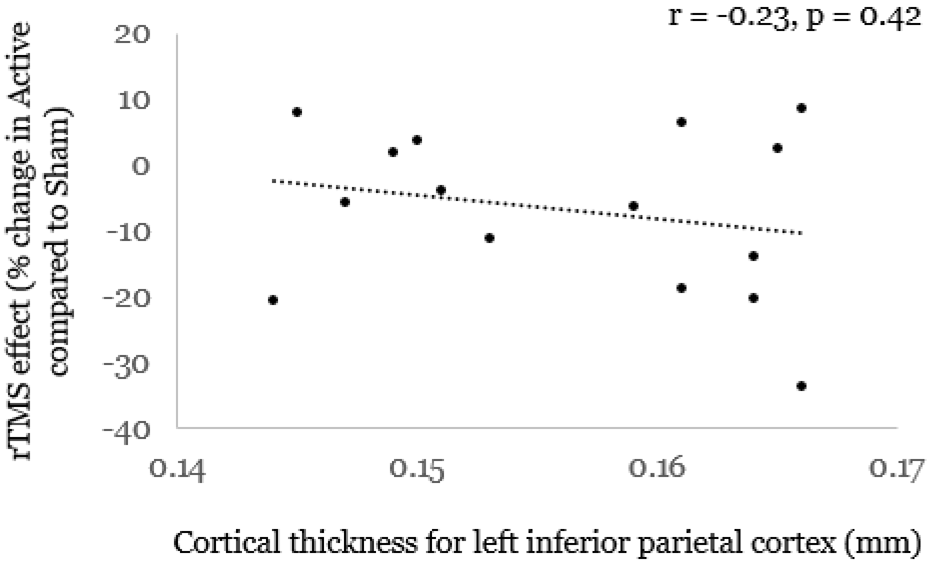


**Figure S5.** Relationship between the cortical thickness at the left parietal target and the rTMS effect obtained in the hardest difficulty level of the DRAT. No significant correlation was found.
